# Extrapolating histone marks across developmental stages, tissues, and species: an enhancer prediction case study

**DOI:** 10.1101/010710

**Authors:** John A. Capra

## Abstract

**Background:** Dynamic activation and inactivation of gene regulatory DNA produce the expression changes that drive the differentiation of cellular lineages. Identifying regulatory regions active during developmental transitions is necessary to understand how the genome specifies complex developmental programs and how these processes are disrupted in disease. Gene regulatory dynamics are mediated by many factors, including the binding of transcription factors (TFs) and the methylation and acetylation of DNA and histones. Genome-wide maps of TF binding and DNA and histone modifications have been generated for many cellular contexts; however, given the diversity and complexity of animal development, these data cover only a small fraction of the cellular and developmental contexts of interest. Thus, there is a need for methods that use existing epigenetic and functional genomics data to analyze the thousands of contexts that remain uncharacterized.

**Results:** To investigate the utility of histone modification data in the analysis of cellular contexts without such data, I evaluated how well genome-wide H3K27ac and H3K4me1 data collected in different developmental stages, tissues, and species were able to predict experimentally validated heart enhancers active at embryonic day 11.5 (E11.5) in mouse. Using a machine-learning approach to integrate the data from different contexts, I found that E11.5 heart enhancers can often be predicted accurately from data from other contexts, and I quantified the contribution of each data source to the predictions. The utility of each dataset correlated with nearness in developmental time and tissue to the target context: data from late developmental stages and adult heart tissues were most informative for predicting E11.5 enhancers, while marks from stem cells and early developmental stages were less informative. Predictions based on data collected in non-heart tissues and in human hearts were better than random, but worse than using data from mouse hearts.

**Conclusions:** The ability of these algorithms to accurately predict developmental enhancers based on data from related, but distinct, cellular contexts suggests that combining computational models with epigenetic data sampled from relevant contexts may be sufficient to enable functional characterization of many cellular contexts of interest.

## Background

Tissue-specific gene regulatory regions are essential for specifying the proper gene expression patterns that drive cellular differentiation and development in animals [17]. However, the recognition of these regulatory regions is challenging. Recently, the ability to assay a range of epigenetic modifications to DNA and histones on a genome-wide scale has improved the ability to interpret the functional potential of non-protein-coding DNA [1]. In particular, several histone modifications, such as monomethylation of the fourth residue of histone H3 (H3K4me1) and acetylation of the 27th residue of H3 (H3K27ac), have been shown to be associated with genomic regions with long range gene regulatory enhancer activity [8, 13, 24].

ENCODE [9], Roadmap Epigenomics [2], and several smaller scale projects have performed thousands of these so-called “functional genomics” assays on hundreds of cell lines and tissue samples. Despite their Herculean efforts, these projects have comprehensively analyzed only a small fraction the modifications in cellular contexts of interest to the scientific community. We are still far from a complete picture of the dynamics of DNA modification and binding across different cells.

Moreover, the initial selection of cellular contexts to characterize by ENCODE was based mainly on practical considerations, such as availability, ease of growth, and high yield. The relevance of functional genomic data from the best characterized cell lines—e.g., the three “Tier 1” ENCODE cell lines: B-lymphocytes (GM12878), embryonic stem cells (H1-hESC), and a leukemia cell line (K562)—to other cellular contexts is unclear due to changes associated with immortalization and the transition to a cancerous state. In addition, these cell lines’ progenitors are developmentally distant from many cells of interest. Indeed, most primary tissues and developmental stages have few data sets available, and these are insufficient to produce a full picture of the functional state of the genome in these cellular contexts.

In this environment, researchers with interests outside of the few well characterized cells are presented with a difficult choice between mapping existing data from other contexts to their own or performing functional genomics analyses in their systems of interest. Furthermore, functional genomics analysis of certain cells may never be possible for technical or ethical reasons, e.g., lack of material or the use of protected tissues. As a result, the mapping of functional genomics data from one context to another is common practice, but the situations in which it is appropriate and the potential pitfalls are not clear. A deeper understanding of the relationships between functional genomics data across contexts is needed to identify the conditions in which mapping across contexts is justified.

Recent work comparing chromatin accessibility and epigenetic modification profiles between pluripotent cells and lineage committed cells has revealed the dynamic nature of these modifications [19, 27, 28, 30]. Embryonic stem cells display more accessible chromatin and potentially active regulatory sequence than differentiated cells [29], and lineage commitment is accompanied by activation of lineage-specific regulatory regions and an overall repression of regions active in embryonic stem cells [25]. The relationships encoded in DNA methylation and chromatin state profiles of different cell types are often sufficient to accurately reconstruct hierarchical relationships based on the anatomical and developmental similarity of cellular lineages [4, 25]. These results suggest that, given the proper models, these relationships could be exploited to “impute” epigenetic information across related cellular contexts to enable functional annotation. Indeed, integrating diverse functional genomics data sets (without regard to their developmental relationships) improves gene regulatory enhancer prediction [10].

In this paper, I evaluate the ability of existing genome-wide H3K4me1 and H3K27ac data sets to identify known gene regulatory enhancers active in a developmentally related, but distinct, context. I focus on heart development in mouse, because this process has both a multi-stage characterization of these two prominent enhancer-associated histone modifications [27] and several hundred experimentally validated enhancers [3, 26]. I introduce a supervised machine learning prediction framework in which I analyze the ability of existing functional genomics data to predict enhancer activity across three dimensions: developmental time within an organism, different tissues within an organism, and corresponding tissues between species. I find that developmental heart enhancers can be predicted very accurately using data from related contexts. Data from all contexts considered, including across species, provide useful information and perform better than random; however, the developmental proximity of a cellular context to the target is correlated with its utility.

## Results

### Preliminaries

My goal was to evaluate the ability of two enhancer-associated histone modifications, H3K4me1 and H3K27ac, collected from different cellular, developmental, and organismal contexts to identify known mouse developmental enhancers (Figure 1). I used H3K4me1 and H3K27ac sites identified via ChIP-Seq on four stages of a directed differentiation of ES cells (E0) to mesoderm (E4) to cardiac precursors (E5.8) to cardiomyocytes (E10) [27]. All other histone mark data I used, including marks from embryonic day 14.5 (E14.5) and eight week old (adult) hearts, were collected by the ENCODE project [9]. Note that the heart data from the first four contexts were collected from a single cell type, while the last two are from full hearts (see Discussion). I took mouse enhancers from the VISTA enhancer browser, a database of experimentally validated DNA sequences (and their genomic locations) that drive gene regulatory enhancer activity at E11.5 in transgenic mice [26]. The mouse genomic regions tested for enhancer activity in VISTA were primarily selected due to evolutionary conservation or association with the P300 transcriptional co-activator protein [3, 15, 26]. The data set consisted of 217 enhancers, 90 of which drove gene expression in the heart. I also considered 88 human DNA sequences shown by VISTA to have heart enhancer activity in transgenic mice in some analyses.

**Figure 1:**
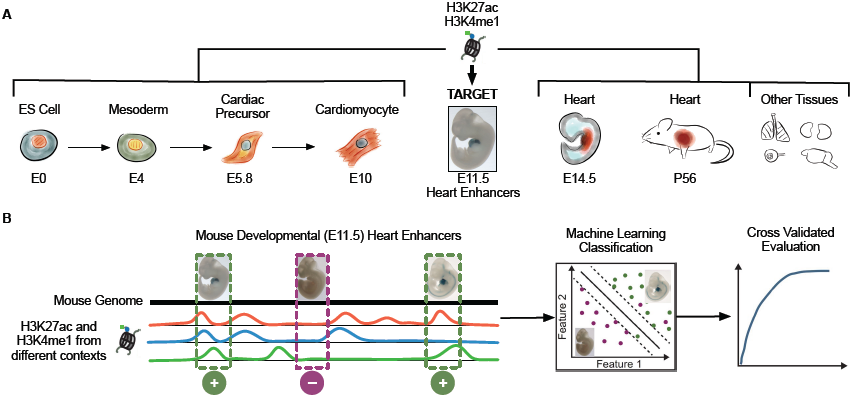
Overview of the data and analyses. **(a)** I collected existing genome-wide maps of two histone marks, H3K4me1 and H3K27ac, from stages of a directed differentiation of mouse embryonic stem (ES) cells into cardiomyocytes, from heart tissues collected from several life stages, and from several other tissues. I evaluated how well these marks, which are associated with enhancer activity, could predict experimentally validated heart enhancers in E11.5 mice (“Target”). **(b)** I took a supervised machine learning approach to this problem by constructing feature vectors for validated enhancers and control regions based on the presence or absence of these histone modifications at their genomic locations. I created classifiers based on different subsets of the data from the cellular contexts given in (a) and evaluated them using cross validation.

I evaluated the ability of the different histone modification data to identify known heart enhancers in a supervised machine learning framework (Figure 1). This type of approach has had great success at identifying enhancers in previous studies [10, 22]. In short, I aimed to classify genomic regions as either positive (having heart enhancer activity at E11.5) or negative (no heart enhancer activity at E11.5) based on the overlap of H3K4me1 and H3K27ac datasets from different cellular, developmental, and species contexts. I refer to these data as “features". In most analyses, validated heart enhancers were the positives. I considered two different sets of negatives: random chromosome and length-matched genomic regions without known enhancer activity and validated enhancers of gene expression in other E11.5 tissues. When random regions were used as negatives, I included 10 matched negatives for each positive. I used six common machine learning algorithms to explore how well these features can predict heart enhancers. I performed five-fold cross validation, in which 20% of the known examples were held out for evaluation of classifiers trained on the remaining data, and evaluated the results by calculating receiver operating characteristic (ROC) curves averaged over the five folds. Unless otherwise stated, I report the results obtained with Random Forests [6], because they performed well and, as described in the Results, the conclusions are robust to the the algorithm used.

### Heart enhancers can be identified accurately using data from different developmental stages

In this section, I evaluate the ability of mouse H3K4me1 and H3K27ac data from different stages of heart development to predict validated mouse heart enhancers at E11.5. I first trained a random forest classifier using all the mouse heart histone mark datasets as features (Figure 1A) to distinguish the heart enhancers from matched regions taken from the genomic background (Methods). This classifier was able to accurately identify E11.5 heart enhancers; the area under the resulting ROC curve (ROC AUC) was 0.96 (Figure 2A).

**Figure 2:**
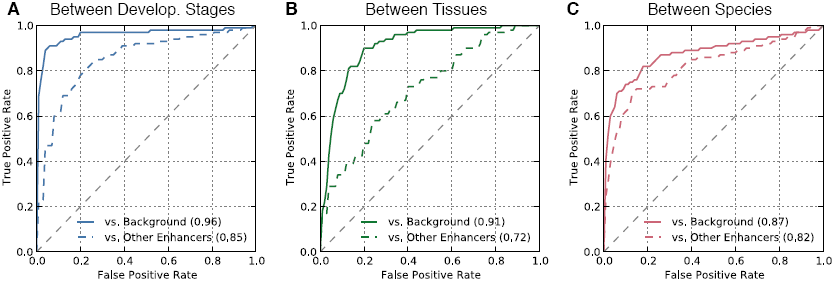
Heart enhancers can be identified accurately using data from different cellular contexts. A random forest classifier was trained to distinguish E11.5 heart enhancers from the genomic background and from enhancers active in other tissues. Each classifier used H3K4me1 and H3K27ac patterns from different sets of cellular contexts as features (Figure 1). (**a**) In five-fold cross validation, the classifiers based on data from other stages of heart development accurately identified E11.5 heart enhancers; these classifiers achieved ROC AUCs of 0.96 vs. the genomic background and 0.85 vs. other enhancers. (**b**) The classifiers that used data from non-heart tissues as features performed well (AUCs of 0.91 and 0.72), but were worse than the developmental-stage-based classifiers. (**c**) When mapped between species, the histone marks from mouse heart development were also able to identify human developmental enhancers better than random (AUCs of 0.87 and 0.82). Note that the results in (**c**) should not be directly compared to the Stage and Tissue results, because they are based on different sets of enhancers. As expected, distinguishing heart enhancers from the genomic background was easier than from non-heart enhancers in each scenario.

Next, I evaluated the ability of classifiers trained using the same features to distinguish the heart enhancers from enhancers active in a diverse set of other tissues (listed in Table S1) at the same developmental stage. As expected from recent results [10], this task proved more challenging, but the random forest classifier still performed well (Figure 2A; ROC AUC=0.85).

### Histone marks from developmental stages nearby the target stage are the most informative

To investigate the contribution of histone marks from different developmental stages to the predictions, I computed the normalized “feature importance” for each feature dataset in the random forest using the Gini impurity metric [6, 20]. For both prediction tasks—heart enhancers vs. background and vs. other enhancers—histone marks from the two stages adjacent to the target stage (E10 and E14.5) had the majority of the importance (Table 1). In both cases, the importance of features decreased monotonically with distance from the target stage. Not surprisingly, given their pluripotent state, marks from embryonic stem cells were the least important.

**Table 1:** Normalized feature importance (Gini impurity) for data from different developmental stages in random forest prediction of E11.5 heart enhancers.

|  | E0 | E4 | E5.8 | E10 | E14.5 | 8 weeks |
| --- | --- | --- | --- | --- | --- | --- |
| Vs. Background | 0.01 | 0.01 | 0.07 | 0.19 | 0.69 | 0.06 |
| Vs. Other Enhancers | 0.05 | 0.07 | 0.17 | 0.37 | 0.23 | 0.11 |

Looking at these data in the simpler terms of the overlap of known enhancers with the different histone marks reveals a similar pattern. I computed the enrichment of each heart histone mark dataset within the E11.5 heart enhancers compared to the background regions. All histone mark datasets showed significant enrichment (*p* < 0.0003 for each; Fisher’s exact test) for overlap with the enhancers (Figure 3; odds ratios >5 for all), but as expected, marks collected from the closest developmental stages to the target (E10 and E14.5) are the most enriched for overlap with the enhancers (odds ratios from 60.5 to 213.9). The embryonic stem cell’s marks showed the least significant enrichment.

**Figure 3:**
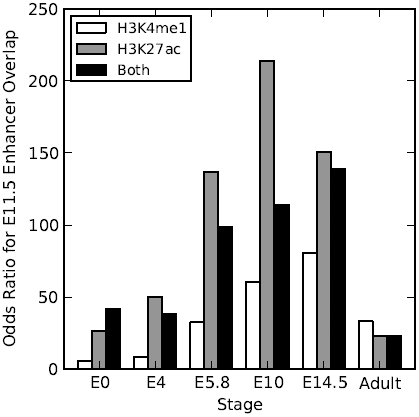
Odds ratios for E11.5 heart enhancer overlap by H3K4me1 and H3K27ac from different cellular contexts. Mouse E11.5 heart enhancers are significantly enriched, compared to matched regions from the genomic background, for H3K4me1 and H3K27ac from all developmental stages listed (*p* 0.01 for all; Fisher’s exact test). As expected, the modifications from contexts flanking E11.5—E10 and E14.5—show the strongest enrichment, while the early stages and adult heart tissue have lower enrichment. Only three E11.5 heart enhancers are not marked by at least one of H3K4me1 or H3K27ac at E14.5.

### Histone marks from different tissues can accurately identify heart enhancers

Next, I evaluated the ability of data from non-cardiac mouse tissues to predict E11.5 heart enhancers. I used H3K27ac and H3K4me1 from 29 different mouse non-cardiac tissues and cell lines collected by the ENCODE project (listed in Table S1) to train random forest classifiers. This classification strategy also performed well at distinguishing heart enhancers from the genomic background (Figure 2B; ROC AUC=0.91); however, it did not achieve the level of accuracy attained with heart data. The non-heart features performed relatively poorly at distinguishing heart enhancers from other types of enhancers (Figure 2B; ROC AUC=0.72). This suggests that these feature data may mark generally active regulatory regions, but do not capture heart-specific patterns.

An H3K4me1 dataset from adult placental tissue received the highest normalized feature importance (10.2%) in the non-heart tissue based classifier of heart enhancers vs. the genomic background. However, I found few clear patterns in the feature importances for these classifiers (Table S2). No tissue dominated the importances, and many datasets from a variety of tissues including liver, limb, embryonic fibroblasts, and brain, had at least 5% of the importance. A largely different set of tissues and marks were found among the most important in the classifier of heart vs. other enhancers.

### Histone marks from mouse heart tissues can accurately identify human heart enhancers

It may never be possible to collect sufficient material for histone mark mapping from some cellular contexts due to ethical or technical reasons. In such cases, using histone marks from model organisms to analyze the target species is appealing. To model this situation, I evaluated the efficacy of using H3K4me1 and H3K27ac collected in mouse hearts to predict known *human* heart enhancers. Studies of the similarity of transcription factor binding [23], methylation [18], and gene expression [5] suggest that this may be feasible due to the considerable similarity in these events in corresponding tissues across distant species.

I mapped validated human heart enhancer sequences to their corresponding locations in the mouse genome, and then used the same random forest strategy to distinguish these regions from the genomic background and other known non-heart human enhancers mapped to mouse. Only 7% of the human heart enhancers overlapped a validated mouse heart enhancer, so they represent a largely independent set of genomic regions to classify. (This should not be taken as an estimate of the number of enhancers conserved between these species, because testing of both the human and mouse sequences for a region was relatively rare.)

The mouse H3K4me1 and H3K27ac heart marks were able to accurately identify the human heart enhancers in both prediction tasks (Figure 2C; ROC AUCs of 0.87 and 0.82); however, they did not perform as well as they did for mouse heart enhancers (Figure 2A). Interestingly, the cross species data were better able to distinguish the human heart enhancers from other types of human enhancer than the marks from mouse non-heart tissues were for mouse heart enhancers (Figure 2B, ROC AUC=0.72). This is consistent with the greater similarity in gene expression in corresponding tissues across species than between different tissues within the same species. However, direct comparison is complicated by the fact that different enhancer regions are being classified in these two analyses.

The human heart enhancers considered here were tested in transgenic mice at E11.5; as a result, the set of human heart enhancers is biased in (at least) two ways. First, they are sufficiently evolutionarily conserved to be mappable between species; 23% of human enhancers did not reliably map to the mouse genome. It is possible that non-conserved human enhancers could be more difficult to predict using mouse data. Second, these regions are active when placed in a mouse. This second bias is unlikely to have a dramatic effect, since the basic transcriptional machinery evolves far more slowly than the regulatory sequences that it acts upon. Indeed, non-conserved enhancer sequences have been shown to maintain activity over much greater distances [11].

### The heart enhancer prediction results are robust to the machine learning algorithm used

Finally, to ensure that the patterns I found in the ability of different histone mark datasets to predict heart enhancers were not specific to a particular classification framework, I repeated all the predictions and evaluations using six common machine learning algorithms: random forests, boosting (AdaBoost), linear support vector machines (SVMs), decision trees, naive Bayes, and *k*-nearest neighbor (KNN) classification. Random forests, AdaBoost, and SVMs all performed similarly well and outperformed the three other approaches (Table 2). No matter the algorithm or overall performance, my general conclusions held: histone marks from diverse contexts can predict heart enhancers better than random, and heart enhancers can be better identified using heart histone mark data than data from other tissues.

**Table 2:** Some classification algorithms perform better than others, but all yield similar conclusions. This table gives ROC AUCs (averaged over five cross-validation folds) for six common algorithms at distinguishing E11.5 heart enhancers from other enhancers based on marks from heart or non-heart tissues.

|  | Heart Features | Non-Heart Features |
| --- | --- | --- |
| Random Forest | 0.85 | 0.72 |
| Linear SVM | 0.84 | 0.73 |
| AdaBoost | 0.82 | 0.70 |
| NaiveBayes | 0.79 | 0.69 |
| Decision Tree | 0.77 | 0.62 |
| KNN ($k = 3$) | 0.74 | 0.66 |

### Histone modifications can be accurately predicted using modifications from other contexts

Thus far, I have focused on predicting enhancers using histone modification data. However, histone modifications are informative about functions beyond enhancer activity, and thus, predicting histone modifications themselves across cellular contexts could have broader utility.

To explore this possibility, I evaluated the ability of mouse histone marks to predict each other. In other words, rather than predicting known enhancers, I held out the histone marks collected from each heart developmental stage in turn and trained a classifier to identify their locations compared to the genomic background using the other marks as features. The locations of both H3K4me1 and H3K27ac at each heart developmental stage were able to be determined more accurately than random guessing (Table 3). However, there was considerable variation in performance. Not surprisingly given their pluripotent state and the broadly active regulatory landscape that accompanies it [25, 29], modifications in E0 cells were by far the most difficult to identify from the available data. Stages in the middle of the differentiation attained the highest scores, likely due to the presence of more temporally flanking modification data.

**Table 3:** ROC AUCs for predicting heart histone modification sites using histone modification data from all other heart developmental contexts.

|  | H3K4me1 | H3K27ac |
| --- | --- | --- |
| E0 | 0.70 | 0.65 |
| E4 | 0.76 | 0.91 |
| E5.8 | 0.88 | 0.92 |
| E10 | 0.94 | 0.95 |
| E14.5 | 0.80 | 0.92 |
| Adult | 0.81 | 0.87 |

## Discussion

These results suggest that there is considerable promise in using functional genomics data across contexts for the common task of identifying putative gene regulatory regions. However, the poor results based on data from early developmental stages indicate that, as expected, there is considerable variability in the utility of different data sets depending on the target context. This underscores the need for methods to highlight the most informative contexts in which to collect new functional genomics data and to “interpolate” across contexts using existing data.

The analyses presented here required the integration of several datasets from different sources, and attributes of these data must be kept in mind while interpreting the results. First, though the heart enhancers validated by VISTA are commonly used to explore attributes of enhancers, they are likely not representative of all developmental heart enhancers. Most heart enhancers in VISTA were selected for analysis based on mammalian evolutionary sequence conservation or the presence of P300 in heart tissue [26]. Thus, it is possible that enhancers that do not have these attributes, e.g., species-specific enhancers, might be harder to identify using histone modifications. Nonetheless, my approach performs very well on this high confidence set of functionally validated enhancers.

To obtain H3K4me1 and H3K27ac from as wide a range of heart developmental contexts as possible, I combined data collected from two different sources. The data from stages prior to the target stage (E0-E10) come from a directed differentiation of cardiomyoctyes, while the data from after the target (E14.5 and adult) were collected from whole hearts. Given that both data types show a clear trend of increasing performance with proximity to the target, I am confident in this trend. However, given this difference in origin, the feature importances assigned to marks from before and after the target stage may not be directly comparable representations of the relevance of these developmental stages to the target.

As more functional genomics data collected over successive developmental stages become available, it will be possible to model the gene regulatory landscapes of different developing tissues. For example, Nord et al. [16] recently collected genome-wide H3K27ac modification profiles from seven developmental stages (starting with E11.5 through adulthood) for mouse heart, liver, and forebrain tissues and found rapid and pervasive turnover in the H3K27ac modification landscape across the development of these tissues. As illustrated in their description of the two phases of liver development, integrating physiological knowledge about the development of different tissues with genome-wide modification profiles will likely be necessary to identify the most informative contexts to assay and to fully characterize developmental enhancer usage.

The analyses presented here take an initial step towards the ultimate goal of understanding how and when functional genomics data should be mapped across contexts. Understanding the dynamics of genome modifications across cell types will be relevant beyond this enhancer prediction case study; genome-wide profiles of TF binding and DNA and histone modification are informative about many possible functions of DNA.

Given the success of these relatively simple models at predicting histone modifications themselves, I anticipate that more explicit models of the dynamics of these features over developmental time in different tissues will enable interpolation of histone marks between different contexts and help identify the most informative contexts to assay. I recently developed a statistical method for modeling DNA methylation dynamics across development that can accurately impute missing methylation data across blood cell differentiation [7]. Similar approaches are likely to enable these analyses for other dynamic modifications and cellular lineages.

## Conclusion

I show that integrating enhancer-associated histone marks from different cellular contexts achieves accurate prediction of heart enhancer activity in a context from which no data were collected. Thus, extrapolating existing functional genomics datasets across developmental, cellular, and species contexts has the potential to enable accurate gene regulatory enhancer prediction in many contexts. More broadly, this suggests the promise of “interpolating” existing functional genomics data to related contexts to complement ongoing experimental efforts to characterize the diversity of mammalian cells.

## Materials and methods

### Data

All analyses were performed using the February 2009 assembly of the human genome (GRCh37/hg19) and the July 2007 assembly of the mouse genome (MGSCv37/mm9). Any data that were not in reference to these builds were mapped over using the liftOver tool from the UCSC Genome Browser’s Kent tools (http://hgdownload.cse.ucsc.edu/admin/jksrc.zip). This tool was also used to map genomic locations between the human and mouse genomes.

Human and mouse enhancer sequences, genomic locations, and expression contexts were downloaded from the VISTA Enhancer Browser [26] on January 6, 2014. This consisted of 217 mouse enhancers, 90 of which had heart activity in E11.5 transgenic mice, and 848 human enhancers, 88 of which had heart activity.

The H3K4me1 and H3K27ac genome-wide profiles were taken from two sources: a recent directed differentiation of mouse embryonic stem cells into cardiomyocytes [27] and mouse heart and non-heart tissues analyzed by the ENCODE project [9]. Each study performed chromatin immunoprecipitation followed by high-throughput sequencing (ChIP-Seq) to identify DNA associated with modified histones. I used the peak calls released by each study. Supplementary Table 1 lists all cellular contexts considered.

### Enhancer Prediction

In this paper, I describe the classification of genomic regions based on their enhancer activity. Different analyses consider different sets of regions as positive and negative training examples as outlined in the Results. However, in general, for each genomic region of interest, a feature vector was created by intersecting the genomic locations of H3K4me1 and H3K27ac peaks from the relevant contexts with its location. The feature vector and the enhancer status were then used to train and evaluate supervised classification algorithms. Chromosome and length matched genomic regions were generated for each set of positives using the randomBed program from the BEDtools suite [21].

I used the python scikit-learn v0.14.1 machine learning module [20] to perform all supervised classification analyses and cross validated evaluations. I used the default scikit-learn implementation of six supervised classification algorithms: Random Forests with 10 trees and maximum depth of 5 [6], linear Support Vector Machines (SVMs) with the *ℓ*^2^ norm and *C* of 0.1, AdaBoost [12] with 50 Decision Trees and learning rate 1, Gaussian Naive Bayes, Decision Trees with maximum depth 5, and K-Nearest Neighbors with *k* of 3. For Random Forests, the scikit-learn implementation combines the probabilistic predictions from each tree, instead of letting each classifier vote for a single class. For Decision Tree and Random Forest classifiers, Gini impurity was used as the metric and to compute feature importance. All other statistical analyses were performed with scipy [14].

## Author’s contributions

JAC performed all analyses and wrote the manuscript.

## Acknowledgements

I am grateful to Katherine Pollard, Dennis Kostka, and members of the Capra Lab for many helpful discussions about this project and related problems. Alex Williams provided original artwork for Figure 1. Alexandra Fish and Corinne Simonti provided comments on the manuscript.

## Additional Files

### Additional file 1 — Table S1

List of H3K3me1 and H3K27ac data sets used in the analyses.

### Additional file 2 — Table S2

Feature importances for Random Forest classifiers using histone marks from non-heart mouse tissues to predict heart enhancers versus the genomic background and versus enhancers of other tissues.

## References

[1] Roger P. Alexander, Gang Fang, Joel Rozowsky, Michael Snyder, and Mark B. Gerstein. Annotating non-coding regions of the genome. Nat Rev Genet, 11(8):559–571, 08 2010.

[2] Bradley E Bernstein, John A Stamatoyannopoulos, Joseph F Costello, Bing Ren, Aleksandar Milosavljevic, Alexander Meissner, Manolis Kellis, Marco A Marra, Arthur L Beaudet, Joseph R Ecker, Peggy J Farnham, Martin Hirst, Eric S Lander, Tarjei S Mikkelsen, and James A Thomson. The nih roadmap epigenomics mapping consortium. Nat Biotech, 28(10):1045–1048, 10 2010.

[3] Matthew J Blow, David J McCulley, Zirong Li, Tao Zhang, Jennifer A Akiyama, Amy Holt, Ingrid Plajzer-Frick, Malak Shoukry, Crystal Wright, Feng Chen, Veena Afzal, James Bristow, Bing Ren, Brian L Black, Edward M Rubin, Axel Visel, and Len A Pennacchio. Chip-seq identification of weakly conserved heart enhancers. Nat Genet, 42(9):806–810, 2010.

[4] Christoph Bock, Isabel Beerman, Wen-Hui Lien, Zachary D. Smith, Hongcang Gu, Patrick Boyle, Andreas Gnirke, Elaine Fuchs, Derrick J. Rossi, and Alexander Meissner. DNA Methylation Dynamics during In Vivo Differentiation of Blood and Skin Stem Cells. Mol Cell, 47(4):633–647, August 2012. ISSN 1097-4164. URL http://dx.doi.org/10.1016/j.molcel.2012.06.019.

[5] David Brawand, Magali Soumillon, Anamaria Necsulea, Philippe Julien, Gábor Csárdi, Patrick Harrigan, Manuela Weier, Angélica Liechti, Ayinuer Aximu-Petri, Martin Kircher, et al. The evolution of gene expression levels in mammalian organs. Nature, 478(7369):343–348, 2011.

[6] Leo Breiman. Random forests. Machine learning, 45(1):5–32, 2001.

[7] John A. Capra and Dennis Kostka. Modeling DNA methylation dynamics with approaches from phylogenetics. Bioinformatics, 30(17):i408–i414, 2014.

[8] Menno P. Creyghton, Albert W. Cheng, G. Grant Welstead, Tristan Kooistra, Bryce W. Carey, Eveline J. Steine, Jacob Hanna, Michael A. Lodato, Garrett M. Frampton, Phillip A. Sharp, Laurie A. Boyer, Richard A. Young, and Rudolf Jaenisch. Histone h3k27ac separates active from poised enhancers and predicts developmental state. Proceedings of the National Academy of Sciences, 107(50):21931–21936, 2010.

[9] Consortium ENCODE Project. An integrated encyclopedia of dna elements in the human genome. Nature, 489(7414):57–74, 2012.

[10] Genevieve D. Erwin, Nir Oksenberg, Rebecca M. Truty, Dennis Kostka, Karl K. Murphy, Nadav Ahituv, Katherine S. Pollard, and John A. Capra. Integrating diverse datasets improves developmental enhancer prediction. PLoS Comput Biol, 10(6):e1003677, 06 2014.

[11] Shannon Fisher, Elizabeth A Grice, Ryan M Vinton, Seneca L Bessling, and Andrew S McCallion. Conservation of ret regulatory function from human to zebrafish without sequence similarity. Science, 312(5771):276–279, 2006.

[12] Yoav Freund and Robert E Schapire. A desicion-theoretic generalization of on-line learning and an application to boosting. In Computational learning theory, pages 23–37. Springer, 1995.

[13] Nathaniel D. Heintzman, Gary C. Hon, R. David Hawkins, Pouya Kheradpour, Alexander Stark, Lindsey F. Harp, Zhen Ye, Leonard K. Lee, Rhona K. Stuart, Christina W. Ching, Keith A. Ching, Jessica E. Antosiewicz-Bourget, Hui Liu, Xinmin Zhang, Roland D. Green, Victor V. Lobanenkov, Ron Stewart, James A. Thomson, Gregory E. Crawford, Manolis Kellis, and Bing Ren. Histone modifications at human enhancers reflect global cell-type-specific gene expression. Nature, 459(7243):108–112, 05 2009.

[14] Eric Jones, Travis Oliphant, Pearu Peterson, et al. SciPy: Open source scientific tools for Python, 2001–. URL www.scipy.org/.

[15] Dalit May, Matthew J Blow, Tommy Kaplan, David J McCulley, Brian C Jensen, Jennifer A Akiyama, Amy Holt, Ingrid Plajzer-Frick, Malak Shoukry, Crystal Wright, Veena Afzal, Paul C Simpson, Edward M Rubin, Brian L Black, James Bristow, Len A Pennacchio, and Axel Visel. Large-scale discovery of enhancers from human heart tissue. Nat Genet, 44(1):89–93, 2012.

[16] Alex S. Nord, Matthew J. Blow, Catia Attanasio, Jennifer A. Akiyama, Amy Holt, Roya Hosseini, Sengthavy Phouanenavong, Ingrid Plajzer-Frick, Malak Shoukry, Veena Afzal, John L. R. Rubenstein, Edward M. Rubin, Len A. Pennacchio, and Axel Visel. Rapid and pervasive changes in genome-wide enhancer usage during mammalian development. Cell, 155(7):1521–1531, 2013.

[17] Chin-Tong Ong and Victor G. Corces. Enhancer function: new insights into the regulation of tissue-specific gene expression. Nat Rev Genet, 12(4):283–293, 04 2011.

[18] Athma A Pai, Jordana T Bell, John C Marioni, Jonathan K Pritchard, and Yoav Gilad. A genome-wide study of dna methylation patterns and gene expression levels in multiple human and chimpanzee tissues. PLoS genetics, 7(2):e1001316, 2011.

[19] Sharon L. Paige, Sean Thomas, Cristi L. Stoick-Cooper, Hao Wang, Lisa Maves, Richard Sandstrom, Lil Pabon, Hans Reinecke, Gabriel Pratt, Gordon Keller, Randall T. Moon, John Stamatoyannopoulos, and Charles E. Murry. A temporal chromatin signature in human embryonic stem cells identifies regulators of cardiac development. Cell, 151(1): 221–232, 2012.

[20] F. Pedregosa, G. Varoquaux, A. Gramfort, V. Michel, B. Thirion, O. Grisel, M. Blondel, P. Prettenhofer, R. Weiss, V. Dubourg, J. Vanderplas, A. Passos, D. Cournapeau, M. Brucher, M. Perrot, and E. Duchesnay. Scikit-learn: Machine learning in Python. Journal of Machine Learning Research, 12:2825–2830, 2011.

[21] Aaron R Quinlan and Ira M Hall. Bedtools: a flexible suite of utilities for comparing genomic features. Bioinformatics, 26(6):841–842, 2010.

[22] Nisha Rajagopal, Wei Xie, Yan Li, Uli Wagner, Wei Wang, John Stamatoyannopoulos, Jason Ernst, Manolis Kellis, and Bing Ren. Rfecs: a random-forest based algorithm for enhancer identification from chromatin state. PLoS computational biology, 9(3): e1002968, 2013.

[23] Dominic Schmidt, Michael D Wilson, Benoit Ballester, Petra C Schwalie, Gordon D Brown, Aileen Marshall, Claudia Kutter, Stephen Watt, Celia P Martinez-Jimenez, Sarah Mackay, et al. Five-vertebrate chip-seq reveals the evolutionary dynamics of transcription factor binding. Science, 328(5981):1036–1040, 2010.

[24] Daria Shlyueva, Gerald Stampfel, and Alexander Stark. Transcriptional enhancers: from properties to genome-wide predictions. Nat Rev Genet, 15(4):272–286, 04 2014.

[25] Andrew B. Stergachis, Shane Neph, Alex Reynolds, Richard Humbert, Brady Miller, Sharon L. Paige, Benjamin Vernot, Jeffrey B. Cheng, Robert E. Thurman, Richard Sandstrom, Eric Haugen, Shelly Heimfeld, Charles E. Murry, Joshua M. Akey, and John A. Stamatoyannopoulos. Developmental Fate and Cellular Maturity Encoded in Human Regulatory DNA Landscapes. Cell, 154(4):888–903, August 2013.

[26] Axel Visel, Simon Minovitsky, Inna Dubchak, and Len A Pennacchio. Vista enhancer browser–a database of tissue-specific human enhancers. Nucleic Acids Res, 35(Database issue):D88–92, 2007.

[27] Joseph A. Wamstad, Jeffrey M. Alexander, Rebecca M. Truty, Avanti Shrikumar, Fugen Li, Kirsten E. Eilertson, Huiming Ding, John N. Wylie, Alexander R. Pico, John A. Capra, Genevieve Erwin, Steven J. Kattman, Gordon M. Keller, Deepak Srivastava, Stuart S. Levine, Katherine S. Pollard, Alisha K. Holloway, Laurie A. Boyer, and Benoit G. Bruneau. Dynamic and coordinated epigenetic regulation of developmental transitions in the cardiac lineage. Cell, 151(1):206–220, 2012.

[28] Wei Xie, Matthew D. Schultz, Ryan Lister, Zhonggang Hou, Nisha Rajagopal, Pradipta Ray, John W. Whitaker, Shulan Tian, R. David Hawkins, Danny Leung, Hongbo Yang, Tao Wang, Ah Y. Lee, Scott A. Swanson, Jiuchun Zhang, Yun Zhu, Audrey Kim, Joseph R. Nery, Mark A. Urich, Samantha Kuan, Chia-an Yen, Sarit Klugman, Pengzhi Yu, Kran Suknuntha, Nicholas E. Propson, Huaming Chen, Lee E. Edsall, Ulrich Wagner, Yan Li, Zhen Ye, Ashwinikumar Kulkarni, Zhenyu Xuan, Wen-Yu Chung, Neil C. Chi, Jessica E. Antosiewicz-Bourget, Igor Slukvin, Ron Stewart, Michael Q. Zhang, Wei Wang, James A. Thomson, Joseph R. Ecker, and Bing Ren. Epigenomic Analysis of Multilineage Differentiation of Human Embryonic Stem Cells. Cell, 153(5):1134–1148, May 2013. ISSN 1097-4172. URL http://dx.doi.org/10.1016/j.cell.2013.04.022.

[29] Jiang Zhu, Mazhar Adli, James Y. Zou, Griet Verstappen, Michael Coyne, Xiaolan Zhang, Timothy Durham, Mohammad Miri, Vikram Deshpande, Philip L. De Jager, David A. Bennett, Joseph A. Houmard, Deborah M. Muoio, Tamer T. Onder, Ray Camahort, Chad A. Cowan, Alexander Meissner, Charles B. Epstein, Noam Shoresh, and Bradley E. Bernstein. Genome-wide Chromatin State Transitions Associated with Developmental and Environmental Cues. Cell, 152(3):642–654, January 2013. ISSN 00928674. URL http://dx.doi.org/10.1016/j.cell.2012.12.033.

[30] Michael J. Ziller, Hongcang Gu, Fabian Muller, Julie Donaghey, Linus T. Y. Tsai, Oliver Kohlbacher, Philip L. De Jager, Evan D. Rosen, David A. Bennett, Bradley E. Bernstein, Andreas Gnirke, and Alexander Meissner. Charting a dynamic DNA methylation landscape of the human genome. Nature, 500(7463):477–481, August 2013. ISSN 0028-0836. URL http://dx.doi.org/10.1038/nature12433.

